# Speech-in-noise perception across the lifespan: A comparative study in Mongolian gerbils and humans

**DOI:** 10.1101/2024.11.06.622262

**Authors:** Carolin Jüchter, Chieh-Ju Chi, Rainer Beutelmann, Georg Martin Klump

## Abstract

Many elderly listeners have difficulties with speech-in-noise perception, even if auditory thresholds in quiet are normal. The mechanisms underlying this compromised speech perception with age are still not understood. For identifying the physiological causes of these age-related speech perception difficulties, an appropriate animal model is needed enabling the use of invasive methods. In a comparative behavioral study, we used young-adult and quiet-aged Mongolian gerbils as well as young and elderly human subjects to investigate the age-related changes in speech-in-noise perception evaluating whether gerbils are an appropriate animal model for the age-related decline in speech-in-noise processing of human listeners.

Gerbils and human subjects had to report a deviant consonant-vowel-consonant combination (CVC) or vowel-consonant-vowel combination (VCV) in a sequence of CVC or VCV standards, respectively. The logatomes were spoken by different speakers and masked by a steady-state speech-shaped noise. Response latencies were measured to generate perceptual maps employing multidimensional scaling, visualizing the subjects’ internal representation of the sounds. By analyzing response latencies for different types of vowels and consonants, we investigated whether aging had similar effects on speech-in-noise perception in gerbils compared to humans. For evaluating peripheral auditory function, auditory brainstem responses and audiograms were measured in gerbils and human subjects, respectively.

We found that the overall phoneme discriminability in gerbils was independent of age, whereas consonant discriminability was declined in humans with age. Response latencies were generally longer in aged than in young gerbils and humans, respectively. Response latency patterns for the discrimination of different vowel or consonant types were different between species, but both gerbils and humans made use of the same articulatory features for phoneme discrimination. The species-specific response latency patterns were mostly unaffected by age across vowel types, while there were differential aging effects on the species-specific response latency patterns of different consonant types.

## Introduction

Speech communication is one of the most important forms of human social interaction. When our ability to communicate is degraded, this puts us at risk of social isolation, cognitive decline and depression [1–3]. Especially elderly people often have to deal with a deterioration in speech processing and understanding, particularly under noisy conditions. This does not only apply to hearing-impaired listeners, but also elderly people with normal audiometric thresholds in quiet suffer from a deteriorated speech perception [4–9]. This so-called hidden hearing loss is one form of presbycusis (age-related hearing loss, ARHL), which can comprise further spectral, temporal and spatial processing deficits [10–13]. Since difficulties in speech perception are a widespread problem in our aging society with major implications for the daily lives of those affected, it is of common interest to elucidate the physiological causes of the age-related deficits in speech-in-noise perception.

Human psychophysical studies have suggested various potential mechanisms underlying speech-in-noise perception deficits. Among others, these include a deterioration in temporal processing ability with age, which would in turn lead to deficits in temporal fine structure (TFS) sensitivity [5,14,15]. Accordingly, temporal processing was found to deteriorate in elderly humans [e.g., 16–19], and a reduced sensitivity to TFS was observed not only in listeners with hearing loss [20], but even in elderly subjects with normal auditory thresholds in quiet [21–23]. Possible physiological causes contributing to an age-related deterioration in TFS sensitivity are deficits in peripheral processing [24] or a decline in central inhibition [25,26]. Other studies reported a reduced ability to use envelope cues for speech recognition in hearing impaired listeners [27,28] or suggested that an age-related imbalance between TFS and envelope cues in noise may result in speech recognition problems [29]. Beyond that, a decline in general cognitive ability involving attention and processing speed as well as a decrease in synchrony of neural firing were hypothesized to contribute to age-related difficulties in speech processing [5,26,30]. Thus, even though the problem is well-known and has been a major focus of research, the physiological causes for the decline in speech processing with age are still under debate.

In order to further investigate the physiological causes of age-related speech-in-noise processing deficits, an appropriate animal model enabling the use of invasive methods is needed. Mongolian gerbils (*Meriones unguiculatus*) have been commonly used for research on speech sound processing [31–36] and ARHL [e.g., 37–44], as well as the interaction between both [45–47]. Gerbils are known for their good hearing sensitivity in the frequency range of human speech [48], and it was demonstrated that vowel and consonant discrimination patterns are similar between young gerbils and young human listeners [31,34]. Moreover, the age-related changes in their peripheral [42,44,49,50] and central [40,51] auditory system are well characterized, and they were proposed to be a well-suited translational model for the understanding of age-related auditory perceptual deficits in human listeners [49]. However, to date no study employing a comprehensive set of speech sounds has investigated how speech-in-noise perception is altered in gerbils with age and how this compares to humans.

Here, we investigated speech-in-noise perception in young and old Mongolian gerbils as well as young and elderly human listeners, employing similar psychophysical paradigms and speech stimuli. We evaluated to what extent gerbils show the same age-related deterioration in speech-in-noise perception as humans and whether they may be an appropriate animal model for the research regarding the underlying physiological causes of the age-related decline in speech-in-noise perception in humans. Young and elderly subjects of both species had to discriminate various consonant-vowel-consonant combinations (CVCs) and vowel-consonant-vowel combinations (VCVs), allowing us to investigate the age-related changes in speech sound discrimination in both gerbils and humans. To investigate the relation between peripheral auditory function and behavioral speech sound discrimination ability we further measured auditory brainstem responses (ABRs) in gerbils and audiograms in human subjects, discussing the potential origins of species-specific differences.

## Materials and methods

### Animals

Thirteen young-adult (4 – 20 months) and ten quiet-aged (33 – 45 months) Mongolian gerbils (*Meriones unguiculatus*) of either sex were used for the experiments. All gerbils were born and raised in the animal facilities of the University of Oldenburg and originated from animals obtained from Charles River laboratories. The animals were housed either alone or in groups of up to three gerbils of the same sex and their cages contained litter, paper towels, cardboard, and paper tubes as cage enrichment. For the period of training and experimental data acquisition, the gerbils were food-deprived in order to increase their motivation during the experiments. Thus, apart from custom-made 10-mg pellets that they received as rewards during the experimental sessions, they were given only restricted amounts of rodent dry food outside of the experiments. The gerbils had unlimited access to water and training took place five days a week. The general condition of the gerbils was checked every day, and their body weights were kept at about 90% of their free-feeding weights. One quiet-aged gerbil died during the data collection period due to a health issue, so that data from this gerbil are missing for the VCV conditions. Data from four of the thirteen young-adult gerbils have been reported previously [34] and small parts of the datasets, that is, data for behavioral discriminations between the vowels /aː/, /eː/ and /iː/ from nine young-adult and all ten quiet-aged gerbils were used for a comparison with data from single auditory nerve fiber (ANF) recordings in a recent study [45]. The care and treatment of the animals as well as all experimental procedures were reviewed and approved by the Niedersächsisches Landesamt für Verbraucherschutz und Lebensmittelsicherheit (LAVES), Lower Saxony, Germany, under permit numbers AZ 33.19-42502-04-15/1990 and AZ 33.19-42502-04-21/3821. All procedures were performed in compliance with the NIH Guide on Methods and Welfare Consideration in Behavioral Research with Animals [52].

### Auditory brainstem response measurements

For evaluating peripheral auditory function, auditory brainstem responses (ABRs) were measured in all gerbils. The animals were anesthetized by an intraperitoneal injection of either a mixture of ketamine (10% ketamine, 71 mg/kg body weight) and xylazine (2% xylazine, 3 mg/kg body weight) diluted in saline (0.9% NaCl) or a mixture of fentanyl (0.005% fentanyl, 0.03 mg/kg body weight), medetomidine (0.1% medetomidine, 0.15 mg/kg body weight) and midazolam (0.5% midazolam, 7.5 mg/kg body weight). Anesthesia was maintained with subcutaneous injections of one-third dose of the initial mixture. Before starting the recordings, all gerbils received a subcutaneous injection of 2 ml saline in order to prevent dehydration, and oxygen supply (0.6 l/min) was provided throughout the measurements. The animals’ body temperature was maintained at approximately 37°C using a feedback-controlled homeothermic blanket (Harvard Apparatus). The ABR recordings were performed inside of a sound-attenuating chamber (IAC 401-A, Industrial Acoustics Company). During the measurements, the head of the gerbil was fixed using a bite bar. Ear bars containing the speakers (IE800, Sennheiser) and calibration microphones (ER7-C, Etymotic Research) were placed in front of the ear canals. The stainless-steel needle recording and reference electrodes were placed subcutaneously at the vertex of the skull and on the midline in the neck, respectively. The electrodes were moistened with saline solution to ensure low impedances. At the beginning of each measurement, the acoustic system was calibrated in situ by measuring the speakers’ frequency characteristics while presenting a sine sweep (0.1 – 22 kHz, logarithmic scaling at 1 octave/s). The speakers’ output during the measurement was then corrected by a minimum phase finite impulse response filter (512^th^ order) that was derived from the impulse responses, leading to flat output levels (±3 dB) for frequencies between 0.3 and 19 kHz. ABRs were recorded in response to clicks (0.2 – 15 kHz, 40 µs duration) with 10-dB level steps (500 repetitions per level). The stimuli were generated using custom-written software in MATLAB (MathWorks), produced at 48 kHz sampling rate by an external audio card (Hammerfall DSP Multiface II, RME), and preamplified (HB7, Tucker Davis Technologies) before presentation. ABRs were amplified (10,000 times) and bandpass filtered (0.3 – 3 kHz) by an amplifier (ISO 80, World Precision Instruments), and digitized using the external audio card (48 kHz sampling rate). Finally, ABR thresholds were defined using custom-written software in MATLAB implementing the approach described in [53], which was visually cross-checked for each threshold and adapted if necessary. All ABR measures reported in Fig 1 are based on the mean threshold, amplitude, or latency of both ears of each animal.

**Fig 1.**
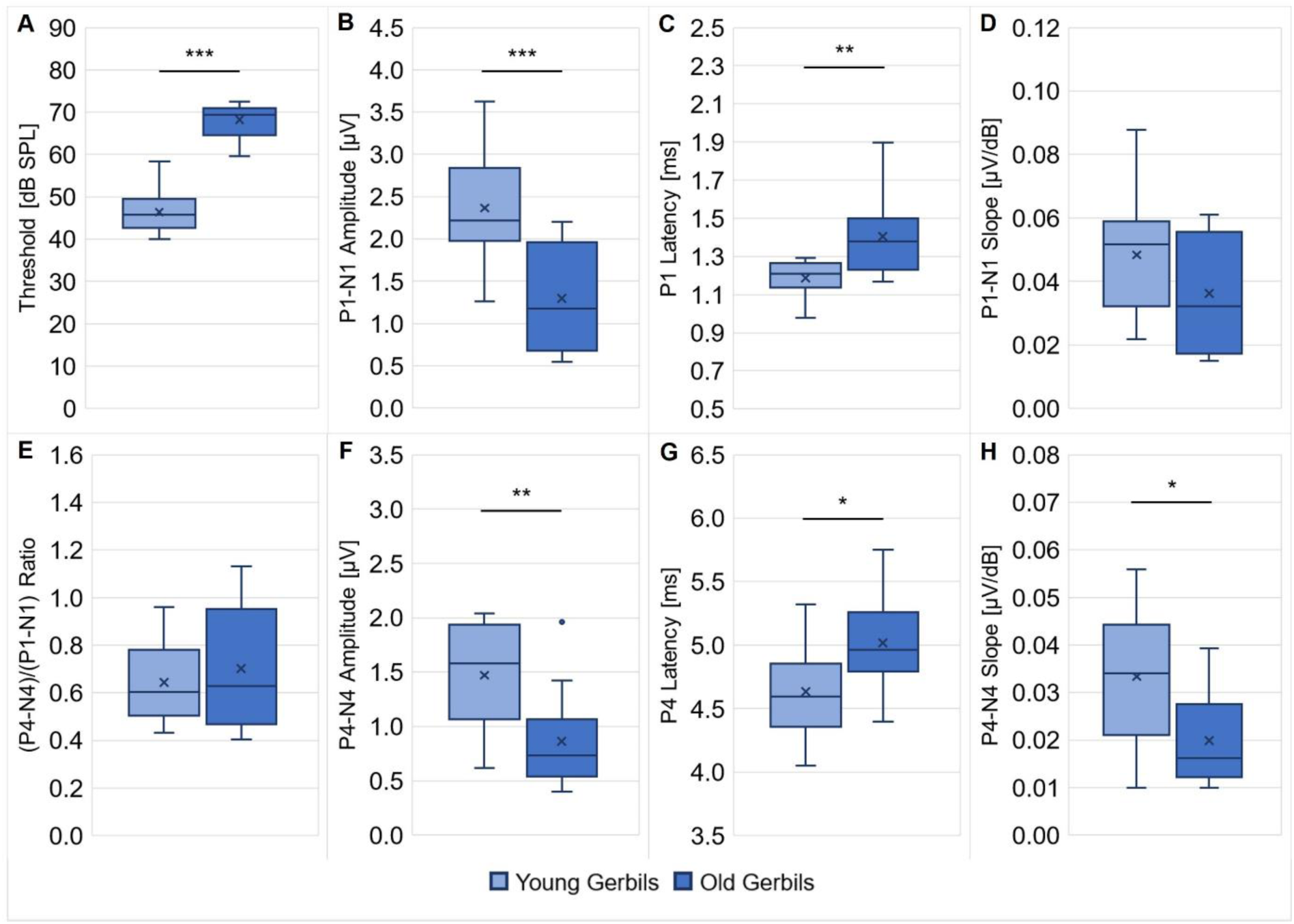
ABRs of young-adult and quiet-aged gerbils. ABRs to clicks were measured in all young-adult and quiet-aged gerbils. ABR thresholds (**A**), P1 latency at 90 dB SPL (**C**) and P4 latency at 90 dB SPL (**G**) were significantly higher/longer in quiet-aged compared to young-adult gerbils. P1-N1 amplitude at 90 dB SPL (**B**), P4-N4 amplitude at 90 dB SPL (**F**) and P4-N4 slope for 60 – 90 dB SPL (**H**) were significantly lower/shallower in quiet-aged gerbils in comparison to young-adult gerbils. No differences between young-adult and quiet-aged gerbils were found in P1-N1 slope for 60 – 90 dB SPL (**D**) and (P4-N4)/(P1-N1) ratio (**E**). These deteriorated ABR measures are clear signs for ARHL in the quiet-aged gerbils. *: *p* < 0.05, **: *p* < 0.01, ***: *p* < 0.001

The measurements indicated a significant difference in ABR thresholds to clicks (unpaired *t*-test: *t*(21) = -11.165, *p* < 0.001; Fig 1A), with on average 22 dB higher thresholds in quiet-aged compared to young-adult gerbils, which is in line with a number of previous studies that reported ABR threshold shifts of typically 15 – 40 dB for old gerbils compared to young gerbils [37–39,45,54,55]. Further, significantly lower P1-N1 amplitudes at 90 dB SPL (unpaired *t*-test: *t*(21) = 3.876, *p* < 0.001; Fig 1B) and significantly longer P1 latencies at 90 dB SPL (unpaired *t*-test: *t*(21) = -3.253, *p* = 0.004; Fig 1C) were found in quiet-aged compared to young-adult gerbils, which corresponds to findings from the literature [51,55,56]. There was no difference in P1-N1 slope for 60 – 90 dB SPL (Fig 1D) and (P4-N4)/(P1-N1) ratio (Fig 1E) between the two age groups. Apart from these changes in ABR threshold and ABR wave I, also age-related deteriorations of ABR wave IV were seen. Matching the observations from previous studies [54–56], quiet-aged gerbils showed significantly lower P4-N4 amplitudes at 90 dB SPL (unpaired *t*-test: *t*(21) = 3.022, *p* = 0.006; Fig 1F), significantly longer P4 latencies at 90 dB SPL (unpaired *t*-test: *t*(21) = -2.472, *p* = 0.022; Fig 1G) and a significantly shallower P4-N4 slope for 60 – 90 dB SPL (Mann-Whitney *U* test: *U* = 28.000, *p* = 0.021; Fig 1H) in comparison to young-adult gerbils. Taken together, these results clearly attest that the peripheral auditory function of the quiet-aged gerbils was declined and that they suffered from ARHL.

### Setup for behavioral experiments

The experimental setup was the same as used in a previous study (for details, see [34]). Briefly, experiments took place in three functionally equivalent setups that were situated in sound-attenuating chambers. In the center of each setup was a custom-built elongated platform with a pedestal in the middle, positioned approximately one meter above the ground. A food bowl connected to an automatic feeder was located at the front end of the platform, facing a loudspeaker used for acoustic stimulation. The movements of the gerbil on the platform and its position on the pedestal were detected by light barriers, and an infrared camera above the platform allowed for additional visual control of the animal during the experiments, which were performed in darkness.

### Behavioral paradigm

The gerbils were trained to perform behavioral experiments as described in detail in [34]. In brief, operant conditioning with food pellets as positive reinforcement was used to train the gerbils to perform an oddball target detection task. During the experiments, the gerbils had to detect a deviating logatome in a sequence of a continuously repeated reference logatome. When the gerbil detected the target logatome, it had to jump off a pedestal to be rewarded with a food pellet. Response latencies and hit rates for the discrimination between all target and reference logatomes were measured. Catch trials, in which the reference logatome did not change, were used in order to determine a false alarm rate as a measure of spontaneous responding.

### Stimuli

The stimulus set selected for the present study comprised 40 CVCs and 36 VCVs originating from the Oldenburg logatome speech corpus (OLLO) [57]. For CVCs, the initial and final consonants were either /b/, /d/, /s/, or /t/ in combination with one of the vowels /a/, /aː/, /ɛ/, /eː/, /ɪ/, /iː/, /ɔ/, /oː/, /ʊ/ or /uː/ in the middle of the logatome. For VCVs, the initial and final vowels were either /a/, /ɪ/ or /ʊ/, combined with one of the medial consonants /b/, /d/, /f/, /g/, /k/, /l/, /m/, /n/, /p/, /s/, /t/, and /v/. The initial and final phonemes within a logatome were always identical (e.g., /bab/ as a CVC or /aba/ as a VCV), and solely the discriminability between logatomes with the same phonetic context was tested so that only a change in the medial phoneme of the logatome had to be detected. Consequently, the gerbils had to discriminate between vowels in the CVC conditions, while VCV conditions were used to test the discriminability of consonants. All logatomes were used both as target and reference logatomes and their order was randomized across sessions and between animals. The logatomes were spoken by two female and two male German speakers and included two tokens per speaker. The speaker and token for each presented reference repetition and the target logatome were randomly chosen. Hence, only a change in the medial phoneme of the logatome, not speaker identity, needed to be reported by the gerbils. Logatomes were presented at 65 dB sound pressure level (SPL) against a continuous noise-masker with speech-like spectral properties (ICRA-1) [58] at 5 dB signal-to-noise ratio (SNR).

### Human data

For the collection of human data on the discrimination of speech sounds, the behavioral paradigm that was used in gerbils was also applied in an adapted version in five young-adult and five elderly human subjects. The young adults (four females, one male) were between 22 and 29 years old, while the elderly human subjects (three females, two males) were aged between 55 and 69 years. All human participants were German native speakers. The experimental procedure for the human subjects was generally similar to that of the gerbils and has been described previously in [34], where also the data of the young-adult human subjects have already been reported together with a subset from the data of nine young-adult gerbils. Different from the gerbil experiments, stimuli were presented to the human subjects via headphones and responses were measured using a touch screen. In the young human subjects, one CVC condition (with the phonetic context /b/) was tested, whereas two CVC conditions (with phonetic contexts /b/ and /s/) were tested in the elderly human subjects. Additionally, all human participants of either age group were tested in two VCV conditions (with phonetic contexts /a/ and /ɪ/). All conditions were tested in the human subjects at a SNR of -7 dB (in contrast to +5 dB SNR for the gerbils), in order to adjust for the previously shown difference in overall sensitivity for human speech sounds between gerbils and humans [34]. The young-adult human subjects participated in the experiments in the course of a student practical course, while the elderly human subjects were recruited for the purpose of the experiment and were paid for their participation. The elderly subjects had already participated in a former unrelated study and were selected because their audiograms showed normal hearing thresholds (below 25 dB hearing level (HL)) in the frequency range most important for speech (0.5 – 8 kHz). This selection was made because we specifically wanted to investigate speech-in-noise problems of elderly human listeners with hidden hearing loss. Accordingly, the pure tone average (PTA, 0.5 – 4 kHz) of the young and elderly human participants did not differ significantly (Mann-Whitney *U* test: *U* = 21.000, *p* = 0.095) and amounted to 1.78 and -3.10 dB HL, respectively. Thus, all human participants were considered to be normal-hearing. However, note that the 8 kHz thresholds of the elderly human subjects were significantly higher than those of the young-adult human subjects (unpaired *t*-test: *t*(8) = -2.319, *p* = 0.049). The experiments were done with the understanding and written consent of each subject following the Code of Ethics of the World Medical Association (Declaration of Helsinki). The procedures were approved by the local ethics committee of the University of Oldenburg.

### Data analysis

Response latencies for the discrimination between all combinations of reference and target logatomes were measured. Confusion matrices filled with the average response latencies for each phoneme comparison of one condition were entered into the multidimensional scaling (MDS) procedure PROXSCAL [59] in SPSS (IBM, version 29). MDS translates the differences in response latencies to perceptual distances in a multidimensional space representing the perceived logatome similarity by spatial proximity. In these perceptual maps, short response latencies are represented by long perceptual distances, since they correspond to a good behavioral discriminability between two logatomes. Long response latencies are reflected by short perceptual distances indicating a poor behavioral discriminability between the logatomes. As a goodness of fit measure for the perceptual maps, the “Dispersion Accounted For” (DAF) was used, which can range from 0 to 1, with high values indicating a better fit. It can be derived from the normalized raw stress (DAF = 1 - normalized raw stress) and provides a measure for the proportion of the sum of the squared disparities (transformed proximities) that is explained by the distances in the MDS solution [60]. The perceptual maps for vowels were arranged in a two-dimensional space, whereas those for consonants were arranged in a three-dimensional space. A higher dimensionality was needed for the consonants in order to reach more than 90% of explained variance in the MDS solutions, leading to similar goodness of fit values for the perceptual maps of vowels and consonants. Adding even more dimensions to the MDS solutions did not lead to a further substantial increase in the amount of explained variance. Apart from that, Spearman’s rank correlations were calculated to compare response latencies between young and old gerbils and human subjects. In addition to response latencies, hit rates and false alarm rates were recorded. For quantifying the subjects’ discrimination ability, the sensitivity-index *d’* was calculated for each subject and CVC or VCV condition, applying the inverse cumulative standard normal distribution function Φ^Φ^ to the mean hit rate *(H)* and mean false alarm rate *(FA)*: *d’* = Φ^Φ^*(H)* – Φ^Φ^*(FA)* [61]. For more details about MDS and the data analysis, see [34].

### Statistics

Statistical analyses were carried out in SPSS (IBM, version 29). Normality of datasets was tested using Shapiro-Wilk tests. To test for age-related differences in various parameters of the gerbils’ ABRs as well as the PTA and thresholds from the audiograms of the human subjects, either two-tailed unpaired *t*-tests or Mann-Whitney *U* tests were used, depending on the distribution of the underlying dataset. For the behavioral data of gerbils and humans, mixed-design ANOVAs were used to test for differences in *d’*-values, response latencies and mean Spearman’s rank correlations of response latencies between different experimental conditions (within-subjects factor) and the two age groups and species (between-subjects factors), respectively. Sphericity of the within-subjects factors were tested with Mauchly’s tests, and the results were adapted with Greenhouse-Geisser corrections when sphericity could not be assumed. Bonferroni-corrected paired *t*-tests were used for post-hoc testing whenever necessary. The threshold for significance (alpha) was set to 0.05 in all statistical tests.

## Results

### Overall behavioral speech sound discrimination ability was independent of age in gerbils, but consonant discriminability for human listeners declined with increasing age

The gerbils’ vowel discrimination ability was tested in four CVC conditions, each with a different consonant as the phonetic context (/b/, /d/, /s/, and /t/). We tested whether the phonetic context had an effect on the overall *d’*-values and response latencies of the young-adult and quiet-aged gerbils during vowel discrimination (S1 Fig). The phonetic context showed no significant effect on either of these behavioral measures. The response latencies were significantly shorter for the young-adult gerbils in comparison to the quiet-aged gerbils (mixed-design ANOVA, factor age group: *F*(1, 21) = 6.017, *p* = 0.023; S1 Fig). However, there were no significant differences in *d’*-values between young-adult and quiet-aged gerbils. No interaction effects between phonetic context and age group were found. Thus, the phonetic context did not affect the gerbils’ overall vowel discrimination ability.

As for the CVC experiments, it was also investigated in multiple VCV conditions whether different vowels as the phonetic context (/a/, /ɪ/, and /ʊ/) affected the gerbils’ overall consonant discrimination ability (S2 Fig). Neither *d’*-values nor response latencies were affected by the different phonetic contexts. As for the CVC conditions, response latencies were significantly shorter in young-adult gerbils in comparison to quiet-aged gerbils (mixed-design ANOVA, factor age group: *F*(1, 20) = 11.583, *p* = 0.003; S2 Fig). However, *d’*-values were not different between the two age groups and no interaction effects between phonetic context and age group were found. These results indicate that the gerbils’ consonant discrimination ability was not affected by the phonetic context of the VCVs.

Since the overall *d’*-values and response latencies of the gerbils were not affected by the phonetic context in the different CVC or VCV conditions, the results from the single conditions were pooled for each age group and species enabling joined analyses of all CVC or VCV conditions, respectively. Thus, we further tested for general differences between CVC and VCV conditions in both gerbils and humans also with respect to their age. The human subjects achieved higher *d’*-values than the gerbils and CVC conditions generally had significantly higher *d’*-values than VCV conditions, while there was no significant difference in *d’*-value by age per se (mixed-design ANOVA, factor logatome type: *F*(1, 28) = 48.375, *p* < 0.001, factor species: *F*(1, 28) = 718.528, *p* < 0.001; Fig 2A). Importantly, all two-way and the three-way interactions turned out to be significant (mixed-design ANOVA, logatome type x age group: *F*(1, 28) = 20.297, *p* < 0.001, logatome type x species: *F*(1, 28) = 8.554, *p* = 0.007, age group x species: *F*(1, 28) = 5.773, *p* = 0.023, logatome type x species x age group: *F*(1, 28) = 14.899, *p* < 0.001; Fig 2A), meaning that each specific combination of logatome type, species and age group had a different influence on the *d’*-value. For example, age only showed a detrimental effect on the *d’*-value of the human participants in VCV conditions, but not in CVC conditions and in neither case for gerbils. Further, all gerbils and the elderly human subjects achieved higher *d’*-values in CVC conditions compared to VCV conditions, but the young human participants showed as high *d’*-values for the VCV conditions as for the CVC conditions. For the response latencies, significant main effects were found for all factors with generally shorter latencies for the discrimination of CVCs compared to VCVs, for young subjects in contrast to old subjects and for humans compared to gerbils (mixed-design ANOVA, factor logatome type: *F*(1, 28) = 35.416, *p* < 0.001, factor age group: *F*(1, 28) = 23.142, *p* < 0.001, factor species: *F*(1, 28) = 123.239, *p* < 0.001; Fig 2B). In contrast to the *d’*-value, there was only one significant interaction effect between logatome type and species on the response latency, indicating that gerbils were significantly slower in VCV conditions compared to CVC conditions, whereas the human subjects were equally fast in both conditions (mixed-design ANOVA, logatome type x species: *F*(1, 28) = 8.310, *p* = 0.007; Fig 2B).

**Fig 2.**
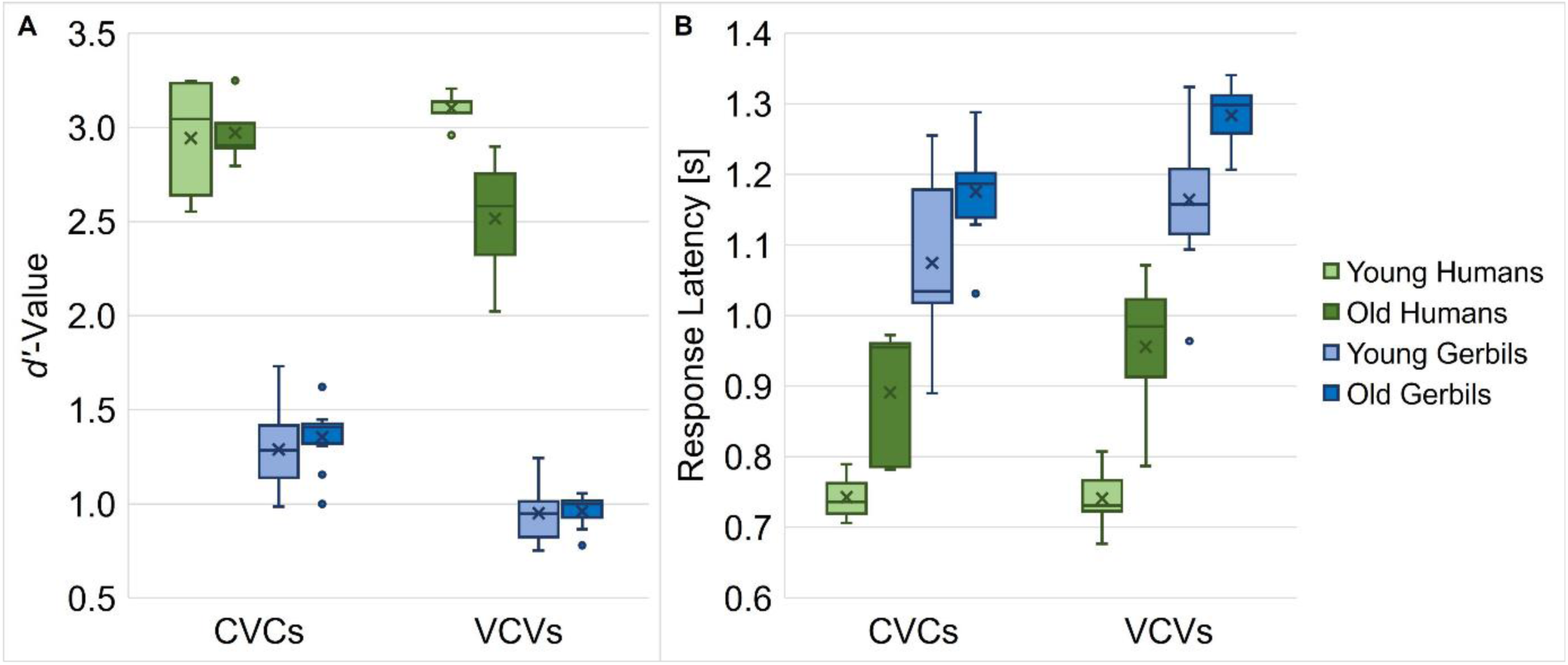
Overall speech sound discrimination ability of young and old gerbils and young and elderly human subjects. Mean *d’*-values (**A**) and response latencies (**B**) of gerbils and humans of both age groups were compared for CVC and VCV conditions. Gerbils showed smaller *d’*-values and longer response latencies than the human subjects, irrespective of the condition. In general, subjects achieved higher *d’*-values in CVC conditions compared to VCV conditions, except for the young human listeners, who were equally sensitive in both conditions. Age had a detrimental effect on the *d’*-values of VCV conditions in human listeners, but not for CVC conditions and in neither case for gerbils. Moreover, age group differences were found for response latencies, with shorter latencies in young subjects compared to old subjects. Further, gerbils were significantly faster in CVC conditions compared to VCV conditions, while human subjects were equally fast in both conditions.

In conclusion, we found differential effects of age on the overall discrimination abilities (as assessed by *d’*-values) of gerbils and humans for vowels and consonants. While humans – in contrast to gerbils – did not show a difference in overall discrimination ability for vowels and consonants in young ages, aging seemed to particularly affect the consonant discrimination ability in humans but not in gerbils. Neither species showed a decline in general vowel discrimination ability with age. Additionally, gerbils were slower in discriminating consonants in comparison to vowels, which was not the case in humans. Generally, humans achieved significantly higher *d’*-values and responded faster than gerbils, which is in line with what we have observed in young humans and a subset of the young gerbils in our previous study [34]. This huge difference in *d’*-value despite the lower SNR for the human participants compared to the gerbils (-7 vs. +5 dB SNR) elucidates the higher difficulty of the experimental task for the gerbils compared to the human subjects. The overall difference in response latency might be due to the differences in the experimental procedure for humans and gerbils, since the human subjects only had to move their finger in order to respond to the target logatomes, while the gerbils had to move their whole body off a pedestal. Further, aging generally led to longer response latencies in both gerbils and humans.

All in all, the overall behavioral speech sound discrimination ability did not decline in quiet-aged gerbils, despite their clear signs of ARHL. In contrast, the elderly human subjects – who were selected for having normal audiometric thresholds – showed a decline specifically in consonant discrimination ability. This indicates that the elderly human subjects were indeed affected by hidden hearing loss, and their decline in consonant discrimination ability contrasts the stable speech-in-noise discrimination performance of the hearing-impaired old gerbils.

### Perceptual maps of vowels and consonants featured similar patterns in gerbils and humans of both age groups

In order to visualize the subjects’ abilities to discriminate between the different vowels and consonants, perceptual maps were generated using MDS. Long distances between two phonemes in the perceptual maps correspond to a good behavioral discriminability, whereas short distances between two phonemes indicate a poor behavioral discriminability.

The two-dimensional perceptual maps for vowels that were generated integrating the data from all CVC conditions of all young or elderly human subjects and young or old gerbils are shown in Fig 3A - D, respectively. Overall arrangement as well as individual locations of the vowels are very similar for the two species and age groups. The ten vowels can be subdivided easily into three separate groups based on their locations in the perceptual maps. These groups reflect the frequency of the second formant (F2) of the vowels: Vowels with high F2 frequencies (/ɛ/, /eː/, /ɪ/ and /iː/) are located on the left side of the perceptual maps, while those with medium F2 frequencies (/a/ and /aː/) are situated close to the middle on the horizontal axis and vowels with low F2 frequencies (/ɔ/, /oː/, /ʊ/ and /uː/) can be found on the right side of the perceptual maps. Thus, the frequency of F2 highly and negatively correlates with Dimension 1 of the perceptual maps (*R*^2^ = 0.862, 0.794, 0.738 and 0.783 for young and old gerbils, and young and elderly human listeners, respectively). An even stronger negative correlation was found for the frequency of the first formant (F1) of the vowels and Dimension 2 (*R*^2^ = 0.947, 0.963, 0.896 and 0.897 for young and old gerbils, and young and elderly human listeners, respectively), which results in a F1 gradient composed of vowels with high F1 frequencies (/a/ and /aː/) being situated in the lower part of the perceptual maps to vowels with medium F1 frequencies (/ɛ/ and /ɔ/) and finally vowels with low F1 frequencies (/eː/, /ɪ/, /iː/, /oː/, /ʊ/ and /uː/) being located in the upper half of the perceptual maps. Note that the vowels within the groups with low F2 frequencies (/ɔ/, /oː/, /ʊ/ and /uː/) and medium F2 frequencies (/a/ and /aː/) were closer together than the vowels within the group with high F2 frequencies (/ɛ/, /eː/, /ɪ/ and /iː/) in the perceptual maps of the human subjects but not in the perceptual maps of the gerbils, meaning that they were perceived as having a higher relative similarity in humans but no in gerbils. However, since the MDS procedure comprises multiple normalization steps, these differences in perceptual distances cannot be transferred to absolute differences in discrimination ability between gerbils and humans.

**Fig 3.**
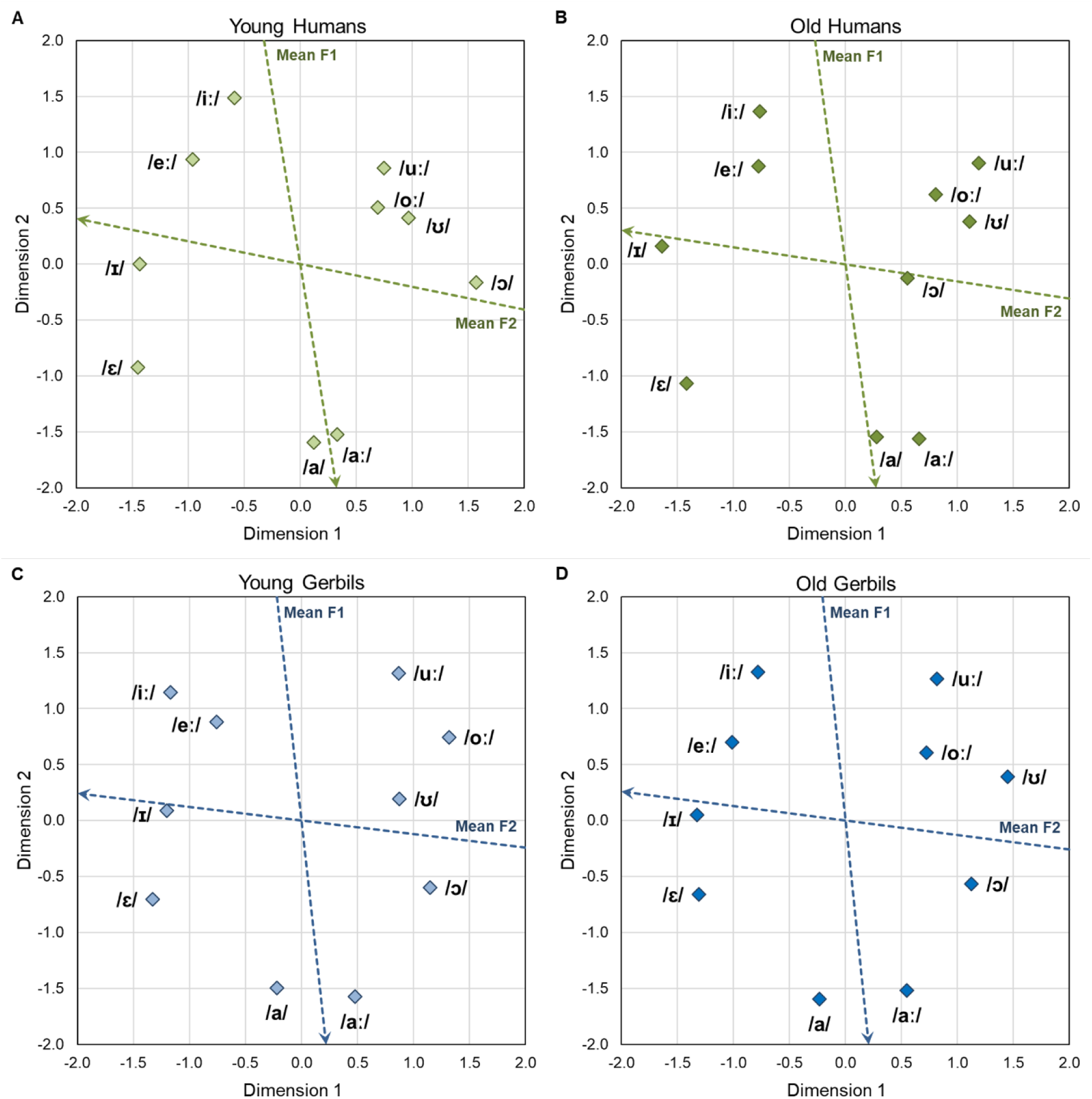
Two-dimensional perceptual maps of gerbils and humans for vowels. Two-dimensional perceptual maps for vowels were generated integrating the data from all CVC conditions of all young (**A**) and elderly (**B**) human listeners as well as young-adult (**C**) and quiet-aged (**D**) gerbils, respectively. Overall arrangement as well as individual locations of the vowels were very similar for all groups. The vowels in the perceptual maps were found to be arranged according to the frequencies of their first two formants. The blue and red dotted arrows show the axes along which the mean frequencies of the first (F1) and the second (F2) formant increase, respectively. The frequencies of F1 and F2 are determined by the tongue height and tongue backness during articulation, respectively.

The overall vowel arrangement found in the perceptual maps of gerbils and humans of both age groups is very similar to the vowel arrangement in the vowel chart for Northern Standard German (see S3 Fig, edited vowel chart for Northern Standard German [62]) which organizes the vowels according to their articulatory features tongue height and tongue backness. The tongue height during articulation determines the frequency of F1, while changes in the tongue backness lead to different F2 frequencies. Consequently, not only humans, but also gerbils of both age groups seem to be able to make use of these human articulatory cues for vowel discrimination. The blue and red arrows in the perceptual maps show property vectors based on a linear regression of the vowel coordinates and the frequencies of F1 and F2, respectively. The DAF values for the MDS solutions of young and old gerbils and young and elderly human listeners amounted to 0.931, 0.934, 0.955 and 0.941, respectively, indicating a very good fit of the perceptual maps to the underlying data. All these findings are consistent with what we have observed previously in a small subset of young-adult gerbils and young human subjects [34]. We found here that these patterns do not only apply to young gerbils and young-adult human subjects, but also to quiet-aged gerbils and elderly human subjects, and that the frequencies of F1 and F2 seem to be the most important cues for the discrimination of vowels in humans and gerbils of all age groups.

Fig 4A - D show the three-dimensional perceptual maps for consonants that were generated integrating the data from all VCV conditions of all young or elderly human listeners and young-adult or quiet-aged gerbils, respectively. All perceptual maps are shown from three different perspectives enabling a better visualization of the three-dimensional arrangement of the consonants. The consonants are marked by different symbols differentiating between various consonant types based on their articulatory features (see S4 Fig, edited consonant chart [63]). The manner of articulation is indicated by color, the place of articulation is marked by shape and the different voicing characteristics can be differentiated by the border of the respective symbol. Depending on the perspective, one can see that the consonants were clustered according to the different characteristics of all of these articulatory features (manner of articulation in the left panels, place of articulation in the medial panels and voicing in the right panels of Fig 4A - D) in gerbils and humans of both age groups. However, different from the vowels with their formant frequencies, there is no such quantifiable correlate for the articulatory features of consonants that could be used for a correlation analysis with the dimensions of the perceptual maps. Also, visually the different characteristics of the articulatory features are not clustered along orthogonal axes so that the articulatory features cannot be assigned one-on-one to the three dimensions of the perceptual maps. Still, a clear clustering of the different articulatory characteristics in the multidimensional space can be seen, which is in line with what we previously found in two-dimensional perceptual maps for consonants in a subset of young-adult gerbils and humans [34] and extends this finding to quiet-aged gerbils and elderly human listeners. The DAF values for the MDS solutions of young and old gerbils and young and elderly humans amounted to 0.943, 0.943, 0.954 and 0.949. Thus, the obtained three-dimensional perceptual maps showed a very good fit to the subjects’ consonant perception reflected by the response latencies.

**Fig 4.**
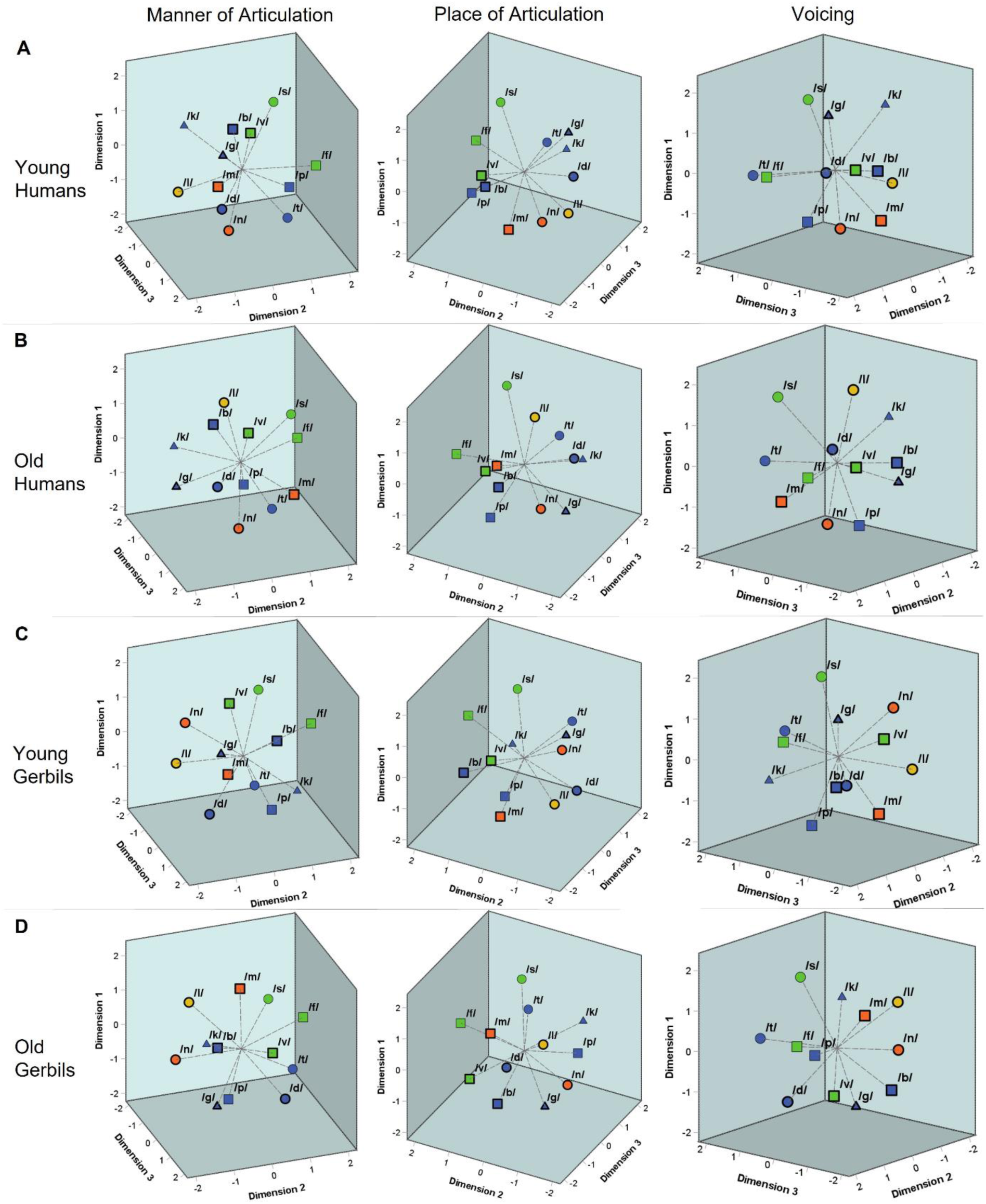
Three-dimensional perceptual maps of gerbils and humans for consonants. Three-dimensional perceptual maps for consonants were generated integrating the data from all VCV conditions of all young (**A**) and elderly (**B**) human listeners as well as young-adult (**C**) and quiet-aged (**D**) gerbils, respectively. All perceptual maps are shown from three different perspectives enabling a better visualization of the three-dimensional arrangement of the consonants. The consonants were found to be clustered according to their articulatory features. The manner of articulation is indicated by color (blue = plosive, green = fricative, orange = nasal, yellow = lateral approximant). The place of articulation is marked by shape (▢ = labial, ○ = coronal, △ = dorsal). The different voicing characteristics can be differentiated by border (**thick border = voiced**, thin border = unvoiced). Depending on the perspective, one can see that the consonants were clustered according to the different characteristics of all of these articulatory features (manner of articulation in the left panels, place of articulation in the central panels and voicing in the right panels) in both age groups of both species.

In a next step, Spearman’s rank correlations for the response latencies of all discriminations between the individual gerbils or humans of each age group were calculated. In order to investigate the inter-individual variability of young and old subjects, the mean correlations of each subject were determined for CVC and VCV conditions (Fig 5). We found significant main effects of the logatome type and the species with generally larger mean Spearman’s rank correlations of the response latencies for CVCs compared to VCVs and for humans compared to gerbils (mixed-design ANOVA, factor logatome type: *F*(1, 28) = 13.580, *p* < 0.001, factor species: *F*(1, 28) = 20.032, *p* < 0.001). Most importantly, significant two-way interactions were observed between the logatome type and the age group as well as the logatome type and the species (mixed-design ANOVA, logatome type x age group: *F*(1, 28) = 15.495, *p* < 0.001, logatome type x species: *F*(1, 28) = 2.623, *p* < 0.001), indicating that there were species- and age-specific differences between the response latency correlations of CVCs and VCVs. Thus, mean Spearman’s rank correlations were significantly higher for VCVs than for CVCs in the human subjects, whereas the correlations were significantly higher for CVCs than for VCVs in gerbils. Further, aging only led to significantly smaller correlations in old subjects compared to young subjects for VCVs, but not for CVCs, meaning that the inter-individual variability for the discrimination of consonants but not for the discrimination of vowels was increased through aging. An overview of the correlations between the mean response latencies for the different age groups of gerbils and humans for vowel and consonant discriminations is shown in the scatterplots in S5 Fig. In summary, there were generally larger inter-individual differences in the response latencies of elderly subjects compared to young subjects in response to consonant discriminations, while there was no effect of aging on the inter-individual variability for vowel discrimination.

**Fig 5.**
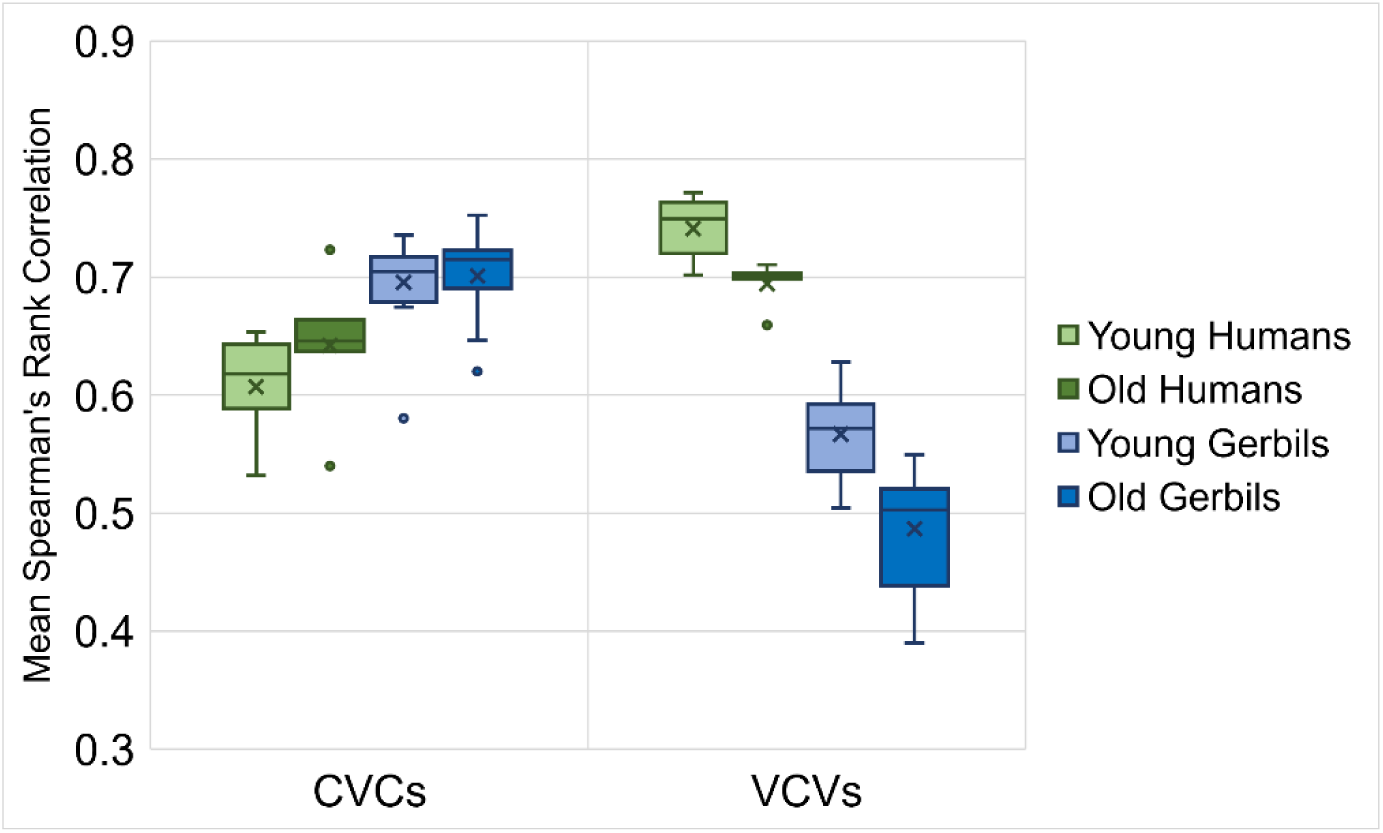
Correlations between response latencies of young and old gerbils and young-adult and elderly human listeners. Spearman’s rank correlations were calculated between the response latencies of all individual gerbils or humans of each age group. Mean correlations between the gerbils were generally higher for CVCs than for VCVs, whereas in humans, correlations were generally higher for VCVs than for CVCs. For CVCs in gerbils and humans, correlations between young subjects were as high as between old subjects. For VCVs, correlations between old subjects were significantly lower than between young subjects in both gerbils and humans.

Taken together, the perceptual maps of vowels and consonants generally showed similar patterns in young and old individuals of gerbils and humans. In both species and independent of the age group, the different types of vowels and consonants determined by their articulatory features were spatially clustered in the perceptual maps, meaning that articulatory similarities also led to a high perceived similarity. However, for consonant discriminations, there were smaller correlations between the response latencies of old subjects compared to young subjects, indicating an age-related increase in inter-individual variability for consonant discrimination.

### Species-specific response latency patterns for discriminating vowel types were mostly unaffected by aging

In order to investigate the discriminability of different types of vowels and consonants in more detail, the response latencies between vowel and consonant pairs from gerbils and humans of both age groups were further evaluated with regard to their articulatory features. For the vowels, the articulatory configurations of both vowels for a specific discrimination were considered with regard to their tongue height (Fig 6A), tongue backness (Fig 6B) and the articulatory features that the two vowels have in common (Fig 6C). To this end, we calculated the mean response latency of each subject for all vowel pairs with a specific combination of articulatory characteristics for the different articulatory features and compared them among each other and between the two species and age groups. In this way, we examined whether there are differences in the response latencies for specific combinations of articulatory features between young-adult and quiet-aged gerbils and young and elderly human listeners. We found significant differences in response latencies for the different combinations of tongue heights and between the species and age groups with shorter response latencies for humans compared to gerbils and young subjects compared to old subjects, respectively (mixed-design ANOVA, factor tongue height: *F*(4.670, 135.419) = 199.790, *p* < 0.001, factor age group: *F*(1, 29) = 19.757, *p* < 0.001, factor species: *F*(1, 29) = 46.965, *p* < 0.001; Fig 6A). Most importantly, a significant three-way interaction effect between tongue height, age and species as well as a significant two-way interaction between tongue height and species were observed (mixed-design ANOVA, tongue height x species: *F*(4.670, 135.419) = 23.942, *p* < 0.001, tongue height x species x age group: *F*(4.670, 135.419) = 2.623, *p* = 0.030; Fig 6A). Thus, specific combinations of tongue height, species and age group had a differential effect on the response latency. For example, the response latencies of young and elderly human subjects were significantly different from each other for all of the different comparisons of tongue heights, while aging only significantly affected the response latencies of gerbils for some of the tongue height comparisons. For all groups, response latencies were longest for the discrimination of two vowels with an open tongue height. Indeed, the response latencies of gerbils and humans for this tongue height comparison were not significantly different in contrast to all other tongue height comparisons.

**Fig 6.**
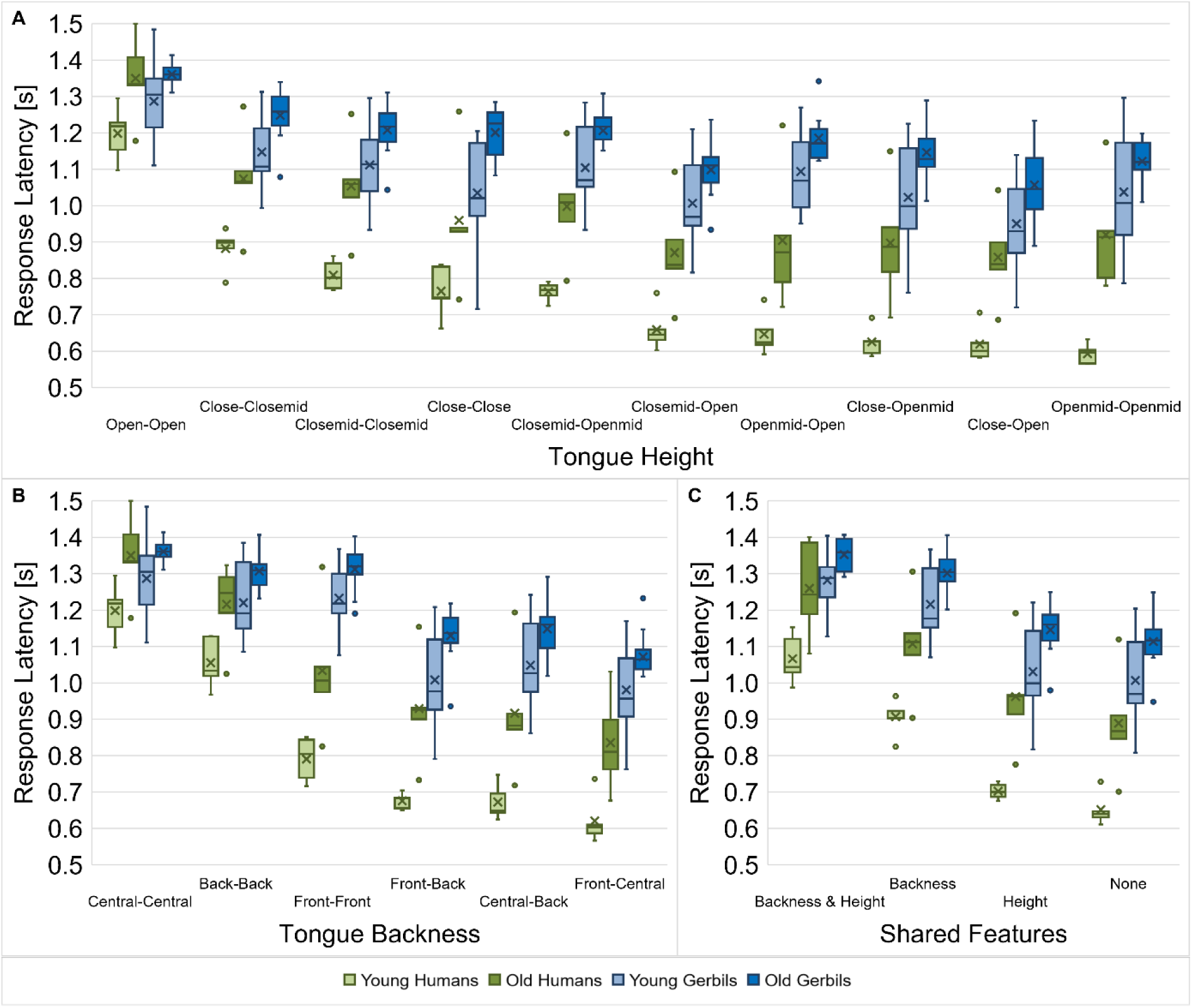
Response latencies for vowel discriminations dependent on the vowel types. Response latencies between vowel pairs from young-adult and quiet-aged gerbils as well as young and elderly human listeners were investigated with regard to their tongue height (**A**), tongue backness (**B**) and shared articulatory features (**C**). Species-specific patterns of the response latencies for the discrimination of different vowel types (dependent on the articulatory features) were found. Aging generally led to longer response latencies in old subjects compared to young subjects. When the vowel comparisons were classified according to the tongue heights during articulation, age did not lead to a consistent increase in response latencies, but it showed differential effects on the species-specific response latency pattern for the different vowel types. Generally, response latencies were longest for vowel discriminations with similar articulatory features, with larger effects of tongue backness than tongue height. The difference in response latency between the easiest and the most difficult discriminations was larger in humans compared to gerbils, meaning that the human listeners were able to benefit more from a lower similarity between vowels than the gerbils.

When the response latencies were classified according to the tongue backness of the vowel comparisons, main effects of tongue backness, species and age group were found (mixed-design ANOVA, factor tongue backness: *F*(2.375, 68.873) = 389.514, *p* < 0.001, factor age group: *F*(1, 29) = 19.028, *p* < 0.001, factor species: *F*(1, 29) = 45.386, *p* < 0.001; Fig 6B). Moreover, there was a significant interaction between tongue backness and species (mixed-design ANOVA, tongue backness x species: *F*(2.375, 68.873) = 49.644, *p* < 0.001; Fig 6B), meaning that the latencies in response to the different combinations of tongue backness changed in a species-specific manner. For instance, note that there is a clear leap in mean response latency from vowel pairs with the same tongue backness (central – central, back – back, and front – front) to vowel pairs with different tongue backness (central – back, front – back, and front – central) in both groups of gerbils, whereas in young and elderly humans the different tongue backness comparisons show more diverse ranges of response latencies. This difference between the species is best visible through the fact that the response latencies for the different combinations of tongue backness are rather similar for the two age groups of each species but show different patterns for humans and gerbils.

Finally, response latencies differed significantly dependent on the articulatory features that the vowel pairs share as well as between the species and age groups (mixed-design ANOVA, factor shared features: *F*(1.704, 49.414) = 456.269, *p* < 0.001, factor age group: *F*(1, 29) = 20.435, *p* < 0.001, factor species: *F*(1, 29) = 46.031, *p* < 0.001; Fig 6C). The response latencies in the pairwise comparisons of the different combinations of shared articulatory features were all differing highly significantly, proving the relevance of the articulatory features for the discriminability of vowels. Especially the same tongue backness during articulation, which determines the F2 frequency, drastically increased the response latency for vowel discriminations in comparison to vowel pairs without any common articulatory features. Tongue height (determining the F1 frequency) had a smaller effect on the response latencies, but still significantly increased the response latency for vowel discriminations. The longest response latencies were seen for vowel pairs that shared both the same tongue backness and tongue height. Additionally, we observed an interaction effect between the shared articulatory features and the species (mixed-design ANOVA, shared features x species: *F*(1.704, 49.414) = 16.987, *p* < 0.001; Fig 6C), indicating that there were species-specific patterns of the response latencies for the different combinations of shared articulatory features.

Altogether, the response latencies for the discrimination of vowels were found to be not only dependent on the species (with generally longer response latencies for gerbils compared to humans) and the vowel type (determined by the articulatory features), but more specifically on the interaction between both. In other words, there were species-specific response latency patterns depending on the articulatory features of the vowels being discriminated. When the vowel comparisons were classified according to the tongue heights during articulation, there was an additional interaction with the age group, meaning that aging had differential effects on these species-specific patterns. This was not the case when the vowel comparisons were classified according to tongue backness or the shared articulatory features, meaning that aging had similar effects on the species-specific response latency patterns in these cases (with consistently longer response latencies in old subjects compared to young subjects). Generally, the results confirm that tongue backness and tongue height are important cues for vowel discriminability in both gerbils and humans, which agrees with what we found previously in a subset of young gerbils and young-adult humans [34]. We observed here that this further applies to quiet-aged gerbils and elderly human listeners. Response latencies were longest for vowel discriminations with similar articulatory features, although differences in tongue height had generally smaller effects than differences in tongue backness. Also, there was generally a larger reduction in response latency for fewer shared articulatory features in humans compared to gerbils, meaning that human listeners could benefit more from the lower similarity of vowels as quantified by the relative difference in response latency compared to the discrimination of vowels with more shared articulatory features.

### Aging differentially affected species-specific response latency patterns for discriminating consonant types

The effect of articulatory features on the discriminability was not only investigated for vowels but also for consonants. Here, the manner of articulation (Fig 7A), voicing (Fig 7B), place of articulation (Fig 7C) and the shared articulatory features (Fig 7D) of the consonant pairs were investigated with regard to differences in their response latencies. As for the vowels, we calculated the mean response latency of each subject for all consonant pairs with a specific combination of articulatory characteristics for the different articulatory features and compared them among each other and between gerbils and human subjects of both age groups. Beside the main effects of age and species, we found that different constellations of manners of articulation led to significantly different response latencies between consonant pairs (mixed-design ANOVA, factor manner of articulation: *F*(3.690, 103.306) = 95.353, *p* < 0.001, factor age group: *F*(1, 28) = 29.517, *p* < 0.001, factor species: *F*(1, 28) = 135.154, *p* < 0.001; Fig 7A). Response latencies were longest for consonant discriminations between two nasal consonants, one nasal consonant and one lateral approximant, or between two plosives, indicating the highest difficulty for the discrimination between these consonant types. In addition to the main effects, there were significant factorial interactions between the manner of articulation and the age group as well as between the manner of articulation and the species (mixed-design ANOVA, manner of articulation x age group: *F*(3.690, 103.306) = 2.998, *p* = 0.025, manner of articulation x species: *F*(3.690, 103.306) = 17.944, *p* < 0.001; Fig 7A). Thus, as for the vowel discriminations, the response latencies showed species-specific patterns depending on the consonants’ manners of articulation. Moreover, also the changes in response latency due to aging were found to be different depending on the consonants’ manners of articulation. The latter is reflected in a smaller age effect on the response latencies for rather difficult discriminations (with comparatively long response latencies) compared to rather easy discriminations (with comparatively short response latencies), e.g., the discrimination of nasal consonants in comparisons to discriminations between consonants with other manners of articulation.

**Fig 7.**
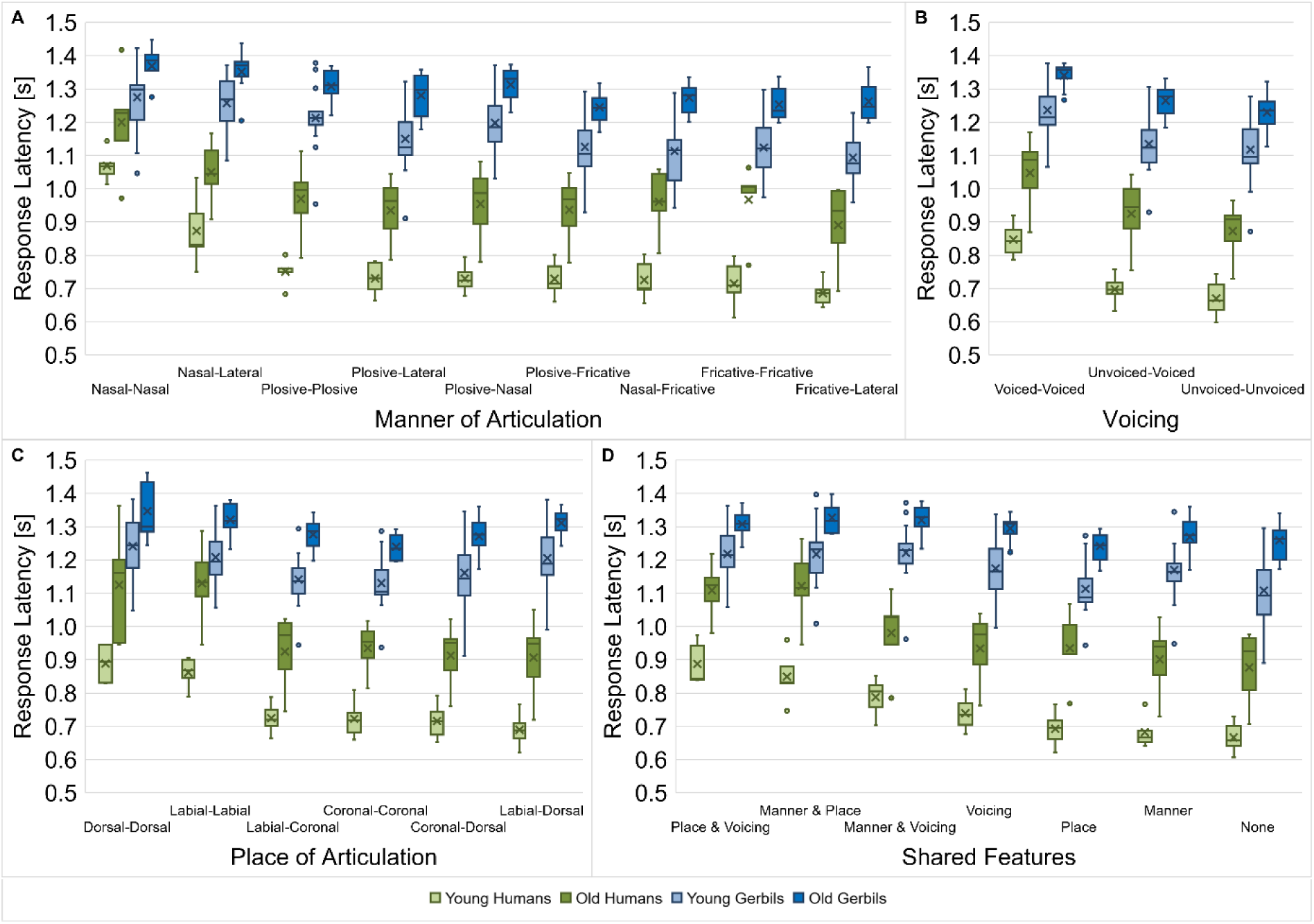
Response latencies for consonant discriminations depending on the consonant Types. Response latencies between consonant pairs from young-adult and quiet-aged gerbils as well as young and elderly human listeners were investigated with regard to their manner of articulation (**A**), voicing (**B**), place of articulation (**C**) and shared articulatory features (**D**). Species-specific response latency patterns were observed for the different types of consonants (dependent on the articulatory features). Age affected the response latencies in a way that old subjects had longer response latencies than young subjects. When the consonant comparisons were classified according to the manner of articulation or the shared articulatory features, age further showed an interaction effect with the different consonant types or/and the species, respectively. Generally, response latencies were longest for consonant discriminations with similar articulatory features. The difference in response latency between the easiest and the most difficult discriminations was larger in young subjects compared to old subjects and in humans compared to gerbils, meaning that they were able to benefit more from a lower similarity between consonants.

When the response latencies were classified according to the voicing of the consonants being discriminated, main effects of voicing, species and age group were found (mixed-design ANOVA, factor voicing: *F*(1.326, 37.136) = 320.273, *p* < 0.001, factor age group: *F*(1, 28) = 26.509, *p* < 0.001, factor species: *F*(1, 28) = 142.153, *p* < 0.001; Fig 7B). Further, we observed an interaction between voicing and species (mixed-design ANOVA, voicing x species: *F*(1.326, 37.136) = 13.905, *p* < 0.001; Fig 7B), with an increasing difference in response latency between gerbils and humans for consonant discriminations with shorter response latencies.

The consonants’ place of articulation also had a significant effect on the response latencies for discrimination as well as the age group and the species (mixed-design ANOVA, factor place of articulation: *F*(1.973, 55.235) = 96.299, *p* < 0.001, factor age group: *F*(1, 28) = 27.661, *p* < 0.001, factor species: *F*(1, 28) = 127.851, *p* < 0.001; Fig 7C). Particularly discriminations between two dorsal or two labial consonants resulted in long response latencies. Again, we found a factorial interaction between the place of articulation and species (mixed-design ANOVA, place of articulation x species: *F*(1.973, 55.235) = 30.300, *p* < 0.001; Fig 7C), resulting in differential response latency patterns of gerbils and humans for the discrimination of consonants with different combinations of places of articulation. These species-specific differences are emphasized by the varying response latency patterns for the different combinations of places of articulation for the two age groups of humans compared to the two age groups of gerbils. For example, note that the response latencies for the discrimination between one labial and one dorsal consonant had the shortest response latencies in both groups of humans, whereas this comparison showed the third longest response latencies in both groups of gerbils.

Lastly, for the shared features, there was again a significant main effect of age and species and a main effect of the shared articulatory features (mixed-design ANOVA, factor shared features: *F*(3.342, 93.588) = 158.067, *p* < 0.001, factor age group: *F*(1, 28) = 28.994, *p* < 0.001, factor species: *F*(1, 28) = 135.452, *p* < 0.001; Fig 7D). Consonant pairs that shared two articulatory features showed the longest response latencies, indicating the worst discriminability. The decrease in discriminability of consonants with an increasing number of shared articulatory features is in line with what we saw previously in a subset of young-adult gerbils and young human listeners [34]. Most importantly, a three-way interaction effect between shared articulatory features, age group and species was found as well as two-way interaction effects between shared articulatory features and age, and shared articulatory features and species (mixed-design ANOVA, shared features x age group: *F*(3.342, 93.588) = 3.045, *p* = 0.028, shared features x species: *F*(3.342, 93.588) = 39.592, *p* < 0.001, shared features x species x age group: *F*(3.342, 93.588) = 3.077, *p* = 0.027; Fig 7D). Thus, specific combinations of shared articulatory features, species and age group had a differential effect on the response latency. Hence, there were response latency patterns that were common to the two age groups of one species, but also patterns that were specific to young or old subjects independent of species. For example, response latencies were comparatively short in young and elderly humans compared to both gerbil groups for consonant discriminations when either the manner of articulation was shared by both consonants or the manner of articulation and the voicing. Then again, discriminations of consonants that share the same manner of articulation and place of articulation showed especially long response latencies in old subjects compared to young subjects, irrespective of the species. Generally, the differences in mean response latencies for the different combinations of shared articulatory features were smaller in old subjects compared to young subjects, indicating that they could not benefit as much from the articulatory differences for consonant discrimination. This effect was even more pronounced in gerbils than in the human subjects.

All in all, the discriminability between consonants was found to be dependent in a species-specific manner on the articulatory features manner of articulation, voicing and place of articulation. When the consonant pairs were classified according to the manner of articulation or the shared articulatory features, the age group further showed an interaction effect on these species-specific response latency patterns. For the consonant pair classifications according to the voicing or the place of articulation, age had a main effect with generally longer response latencies in old subjects compared to young subjects. All subjects showed the longest response latencies – corresponding to the worst discrimination ability – for consonants that have two articulatory features in common. However, old subjects were not able to benefit as much from a lower similarity of consonants for their discrimination ability as young subjects.

## Discussion

In the present study, a behavioral paradigm was used to investigate and compare the age-related changes in the ability for speech sound discrimination in gerbils and humans. In the following, we will discuss differences in speech-in-noise perception between gerbils and humans and evaluate whether gerbils are an appropriate animal model for the known age-related deteriorations in speech-in-noise processing in elderly human listeners.

The overall speech-sound discrimination ability – as assessed by mean *d’*-values – was significantly lower in gerbils compared to humans (Fig 2A) and gerbils needed a higher SNR than humans in order to successfully discriminate vowels and consonants. This is in line with previous reports [31,34] and may be explained by species-specific differences in general psychoacoustic capacities, such as a lower frequency selectivity in gerbils reflected in wider auditory filter bandwidths [64,65], and large differences in familiarization and overall importance of speech sounds for gerbils and humans. Apart from that, aging led to generally longer response latencies in both gerbils and humans (Fig 2B). This finding is consistent with previous studies that also reported longer response latencies – even independent of hearing loss – in old subjects compared to young subjects both in gerbils [41,66] as well as in human listeners [67]. This effect may be observed especially when a motor response is required in response to stimuli with unpredictable timing [68,69], as it is in our behavioral paradigm. Further, an increase in listening effort as it has been previously observed in elderly listeners and listeners with hearing loss [70,71] might contribute to the increase in response latencies, since subjective listening effort has been shown to correlate with response times [72]. Thus, the observed overall increase in response latency in old gerbils and elderly human subjects was unlikely a result of effects related to cochlear aging in particular, but rather could reflect a general age-related cognitive and/or motor decline.

In a profound analysis regarding potential differences between various types of vowels (Fig 6) and consonants (Fig 7) with different articulatory features, we observed that the response latencies for the discrimination of vowels and consonants did not only differ dependent on the articulatory features, but there were also species-specific and age-related differences that showed interaction effects with the articulatory features. The age-related changes in overall vowel and consonant discrimination differed between gerbils and humans. On the one hand, we found that the overall behavioral speech sound discrimination ability in quiet-aged gerbils compared to young-adult gerbils as assessed by *d’*-values was not reduced although they suffered from ARHL. On the other hand, a significant decline in overall consonant discrimination ability was observed in the elderly human subjects, despite their hearing thresholds being similar to those of the young-adult human subjects. Possible explanations for these observations will be discussed below.

### Why is vowel discrimination spared from age-related decline in speech-in-noise processing in gerbils and humans?

The perceptual maps for vowels generated from the response latencies in the behavioral paradigm showed a very high similarity (Fig 3), suggesting that the perception of vowels and the cues used for vowel discrimination are largely identical in young and elderly subjects of gerbils and humans. Further, even though there was an overall main effect of age on the response latencies in both species, no decline in discrimination sensitivity between young and old subjects was observed for vowel discriminations in gerbils and humans (Fig 2A). Also, the correlations of the response latencies of young and old subjects were high and similar for both species (Fig 5), meaning that the overall vowel discrimination ability of gerbils and humans was largely unaltered by aging.

Our results correspond to findings from previous studies reporting that age-related problems in speech perception are less prevalent for vowels than for consonants and that consonants play a more important role in the age-related degradation of speech intelligibility in humans [7,72,73]. Further, identification errors between vowels were found to be largely similar for young and elderly human subjects [74,75], corresponding to the high similarity between the perceptual maps of our young and elderly subjects. Even a considerable hearing loss in human listeners showed only minimal effects on vowel identification [76]. Also for gerbils, the results correspond to findings from a previous study, in which old gerbils were found to have similar discrimination thresholds for the vowel pair /ɪ/-/i/ as young gerbils [47]. Likewise, another study that determined difference limens within speech continua in gerbils showed that age did not affect the behavioral vowel discrimination performance [46]. Thus, there is increasing evidence that aging does not lead to a decline in the behavioral vowel discrimination ability in both gerbils and humans. Interestingly, when comparing the behavioral data for the discrimination of a small subset of vowels (/aː/, /eː/ and /iː/) in gerbils with data from recordings of single ANFs, it was observed that the discrimination based on the temporal responses of ANFs was even improved in old gerbils [45]. The improved temporal coding of ANFs for vowels in old gerbils could be (at least partly) explained by their elevated thresholds, because stimulating closer to threshold (which applies for a fixed stimulus level of 65 dB SPL for both young and old gerbils) resulted in enhanced envelope encoding in the ANFs of quiet-aged compared to young-adult gerbils [45,77]. Since the distinct formant frequency pattern (especially F1 and F2) resulting in a typical temporal waveform is most important for vowel discrimination [78–80], the enhanced envelope encoding in the ANFs of old gerbils might be especially beneficial for discriminating vowels. Consequently, the results suggest that there are other age-related deteriorating processes probably in the central auditory system reducing the sensitivity of quiet-aged gerbils in a way that subsequently the behavioral discrimination ability matches that of young-adult gerbils [45].

A potential candidate for such a central age-related deteriorating process is the decrease in temporal selectivity and heterogeneity of temporal responses between neurons [40,81] due to an age-related decrease in inhibition at multiple stages along the auditory pathway [82–89], which possibly results in a larger redundancy of responses to speech sounds [45]. Thus, a decline in central inhibition has been suggested to be causal for the impaired temporal auditory processing in elderly listeners [89].

Another reason why vowel discrimination is spared from age-related deteriorations in contrast to consonant discrimination might be that vowels comprise a dominant low-frequency formant structure, while age-related hearing loss predominantly affects high frequency regions [42]. Thus, particularly high frequency speech cues but not low frequency cues might be affected by age-related hearing loss, which are more prevalent in consonants than in vowels [90,91].

### What makes humans more vulnerable to age-related declines in consonant perception compared to gerbils?

Unlike the unchanged overall discriminability of vowels in old gerbils and elderly human subjects, we found some age-related differences in the discrimination ability for consonants. The results showed a significant decrease in overall consonant discrimination ability in elderly human listeners compared to young human listeners (Fig 2A). Deficits in consonant recognition and discrimination (in particular in noise) are already known from elderly human listeners, even when they show otherwise normal hearing abilities [5,92], as the human subjects in the present study did. An additional hearing impairment can still lead to even more pronounced difficulties in consonant perception compared to elderly listeners with normal hearing thresholds [93,94]. However, the audiometric hearing threshold can only explain small parts of the variance in consonant recognition in noise [95]. Apart from that, the overall organization of consonants in the perceptual maps generally showed similar patterns for young and old gerbils and young-adult and elderly human subjects (Fig 4), indicating that the cues used for consonant discrimination were generally similar in gerbils and humans of both age groups. This is in line with what was observed previously in humans, where the presence and type of hearing loss in elderly humans affected the overall performance, but not the specific consonant error patterns [92,93,96]. In other words, even though the overall performance decreased with age in elderly humans, consonant discriminations that were most difficult for young listeners were also most difficult for elderly listeners and easy discriminations were perceived as such for listeners of all ages.

In addition to the decrease in overall consonant discrimination ability in elderly human listeners, we observed significantly lower correlations of the response latencies for consonant discriminations between the elderly subjects compared to the young subjects in both species (Fig 5). Hence, there was a higher inter-individual variability in consonant discrimination in elderly subjects compared to young subjects. An increase in inter-individual variability as well as an overall decrease in sensitivity might be caused by a decrease in central auditory temporal precision that becomes increasingly important for complex stimuli as consonants. Not only that the precise temporal representation of neural responses is generally needed for capturing the fast changing acoustic transitions that characterize consonants [97], but also that TFS sensitivity was found to be the best single predictor for modelling consonant identification [5].

As discussed earlier, aging and ARHL involve a deterioration in temporal processing that may result from a decline in central inhibition. This age-related change may be especially important for the differentiation of temporally complex sounds such as consonants. Depending on the severity of the ARHL of the individual subject, the temporal processing deficit may be more or less strong. Consequently, there would be a higher inter-individual variability in temporal processing abilities in old subjects compared to young subjects. This would be in line with a previous study that found an impaired temporal resolution in a behavioral gap detection task for only some of the tested old gerbils, while the other old gerbils showed no such age-related deterioration [41]. A decreased auditory temporal precision has also been observed in aged rats [98] and was hypothesized to be linked to higher inter-individual variability in old animals [26]. Indeed, also in elderly human listeners a higher inter-individual variability [99–102] and increased gap detection thresholds [10] as well as highly variable phoneme boundaries in syllable identification [74] have been observed. Additionally, the identification of syllables with different consonants in old human subjects depended more strongly on stimulus level than in young human subjects [103]. Thus, age-related changes in the central auditory system, such as a decrease in the precise temporal representation of complex sounds, may contribute to the larger variability and an overall decline in consonant discrimination in elderly listeners [26].

A possible reason for the differences in the effect of aging on the overall consonant discrimination ability in gerbils and humans might be that aging and ARHL in humans often coincides to some degree with (cumulative) noise-induced hearing loss (NIHL) due to a repeated exposure to loud noises over the life time [4]. In contrast, it is unlikely that the old gerbils that were used in the present study were affected by NIHL, since they were raised and kept under controlled quiet conditions. Consequently, elderly human subjects with a mixture of ARHL and NIHL might show larger declines in speech-in-noise perception in comparison to old gerbils that were not exposed to loud noise. Indeed, it has been previously observed that elderly human listeners who also show signs for NIHL tend to exhibit threshold shifts in high frequency areas [104]. Since consonants in contrast to vowels comprise more high-frequency components, a mixture of ARHL and NIHL may particularly affect the perception and discrimination of consonants. Indeed, the 8 kHz thresholds in the audiograms of the human subjects in the present study were significantly higher in the elderly humans compared to the young-adult humans, and they were significantly and negatively correlated with the mean hit rate of the VCV conditions (Spearman’s rank correlation: *r*_s_(10) = -0.636, *p* = 0.048). Thus, the increased high-frequency threshold may in fact contribute to the lower hit rate of the elderly human subjects for consonant discriminations compared to the young-adult human subjects.

However, not only NIHL but also ARHL is reflected mostly in threshold shifts in high frequency regions [42]. Thus, the elevated threshold in the click-ABRs of the old gerbils (Fig 1A) probably also arises (at least partly) from threshold shifts at high frequencies [37,54]. It is therefore unlikely that the differences in age-related changes in consonant discrimination between elderly humans and gerbils can be explained solely by different patterns of high-frequency hearing loss, but that there are other species-specific differences that lead to these disparities.

## Conclusion

All in all, we saw that there are many similarities between gerbils and humans in speech-in-noise perception, despite the large difference in overall sensitivity for human speech sound discrimination as assessed by *d’*-values. The similarity of the perceptual maps of both species suggests that they make use of the same articulatory cues for phoneme discrimination, even if the weighting of these cues might differ to some extent between the species for the different types of vowels and consonants. In general, aging led to an increase in the response latencies in both species. These longer response latencies did not translate into a reduced speech sound discrimination ability per se. However, aging affected consonant processing in elderly human subjects. In contrast, vowel perception in both species as well as consonant perception in gerbils were left mostly unaltered by aging. Consonant discrimination might be more vulnerable to age-related declines than vowel discrimination since aging is accompanied by declines in auditory temporal precision, which might be especially important for sounds with a temporally complex structure as consonants. Further, ARHL mostly affects high frequency regions, which are more important for discriminating consonants than vowels. An elevation of high-frequency thresholds may be particularly observed when ARHL coincides with some degree of NIHL – a state that can be observed regularly in elderly human subjects, but that is unlikely in quiet-aged gerbils. However, there might also be other species-specific differences that led to the differences in age effects on the consonant discrimination ability between humans and gerbils.

Taken together, gerbils might be a good model for the general mechanisms of vowel discrimination in humans of all age groups, provided that their differences in overall sensitivity and species-specific discrimination patterns are thoroughly considered. The same applies for consonant discrimination in young normal-hearing subjects, however, since old gerbils did not show the same deteriorations in consonant discrimination ability as elderly human listeners, they might not be an appropriate model for the research regarding the underlying physiological causes of the age-related decline in consonant perception in humans.

## Acknowledgements

We thank Melissa Jäger and Swantje Preller for their contribution to collection of the gerbil data and their care for the animals during the data collection period. Further, we thank Jessica Enter for her contribution to the collection of the human data. Special thanks are dedicated to the five elderly human subjects that took part in the study and to Dawid Fandrich, Franziska Berger, Laura-Janine Döring, Melissa Jäger and Nadine Dyszkant for their participation in the study as part of a student course.

## Supporting information

**S1 Fig.**
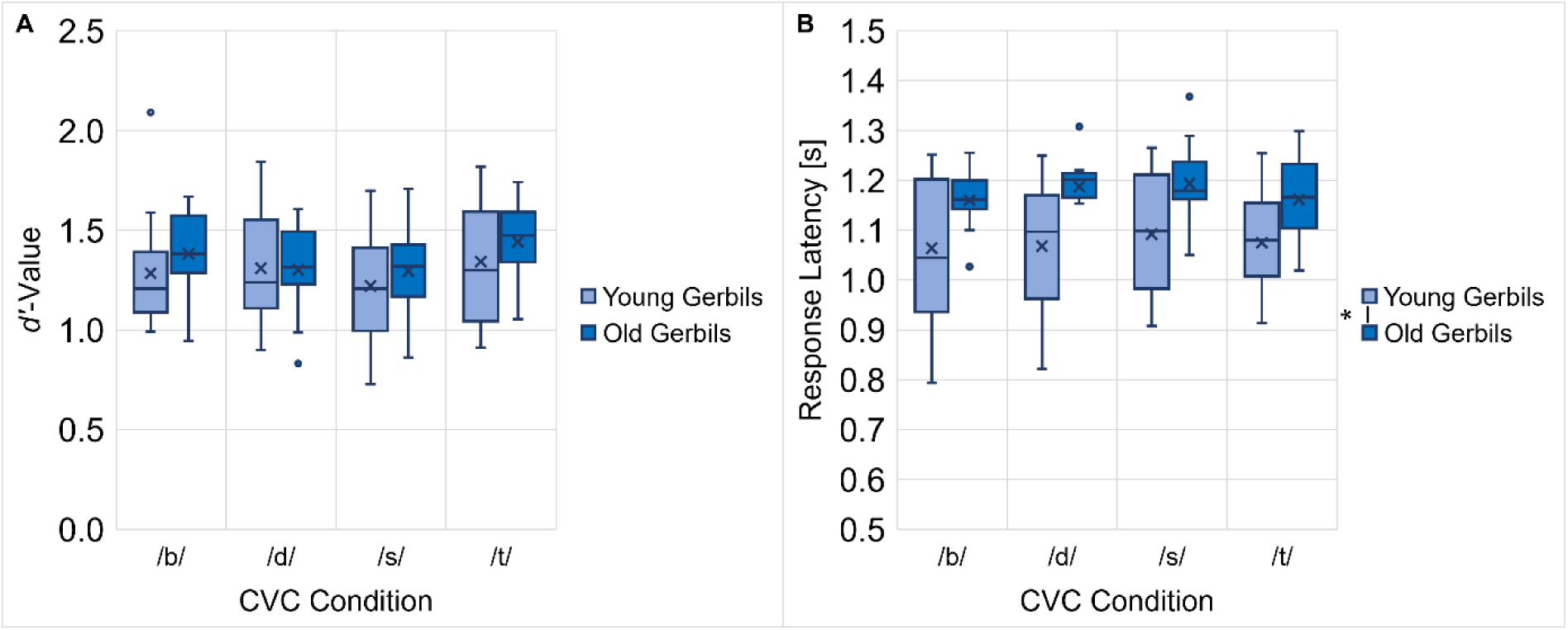
Influence of different flanking consonants on vowel discrimination in gerbils. *: *p* < 0.05

**S2 Fig.**
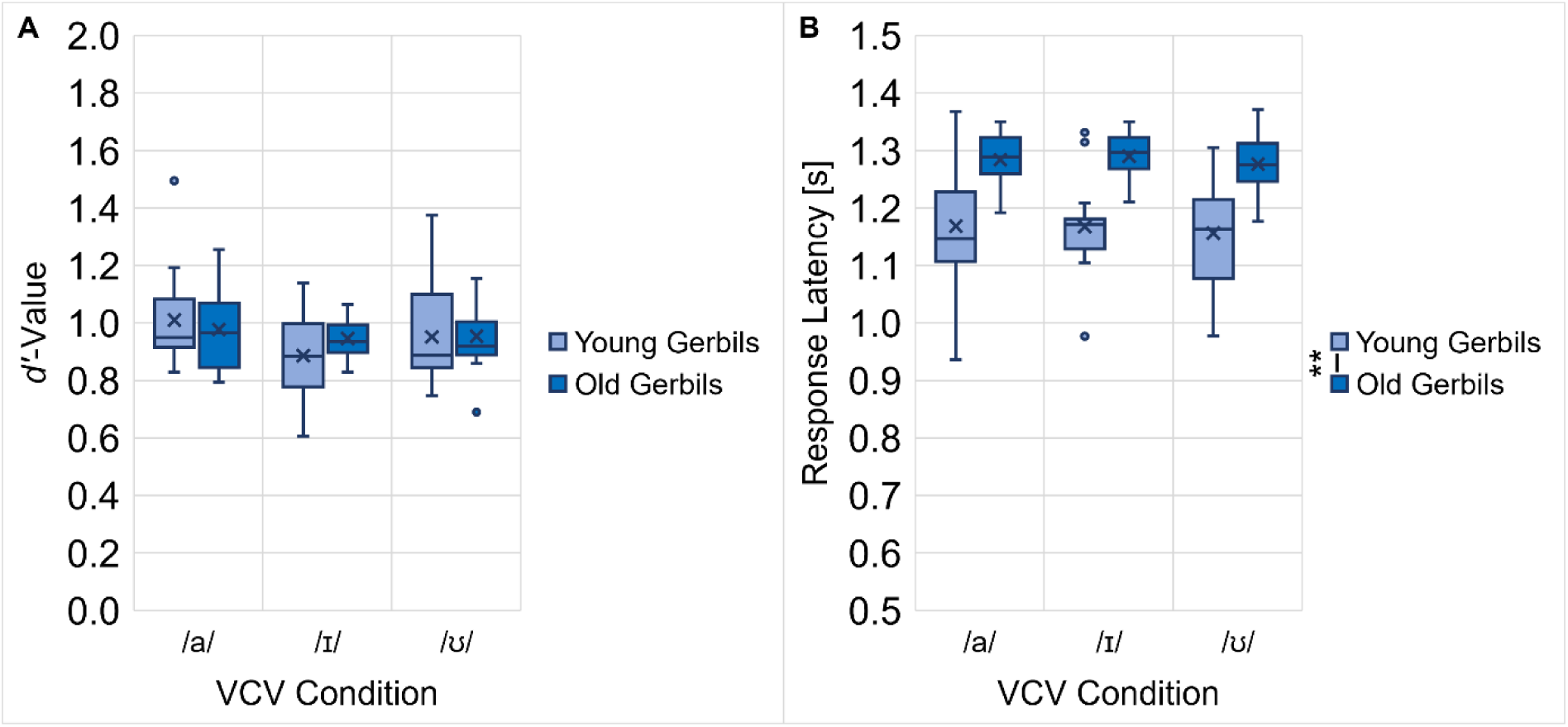
Influence of different flanking vowels on consonant discrimination in gerbils. **: *p* < 0.01

**S3 Fig.**
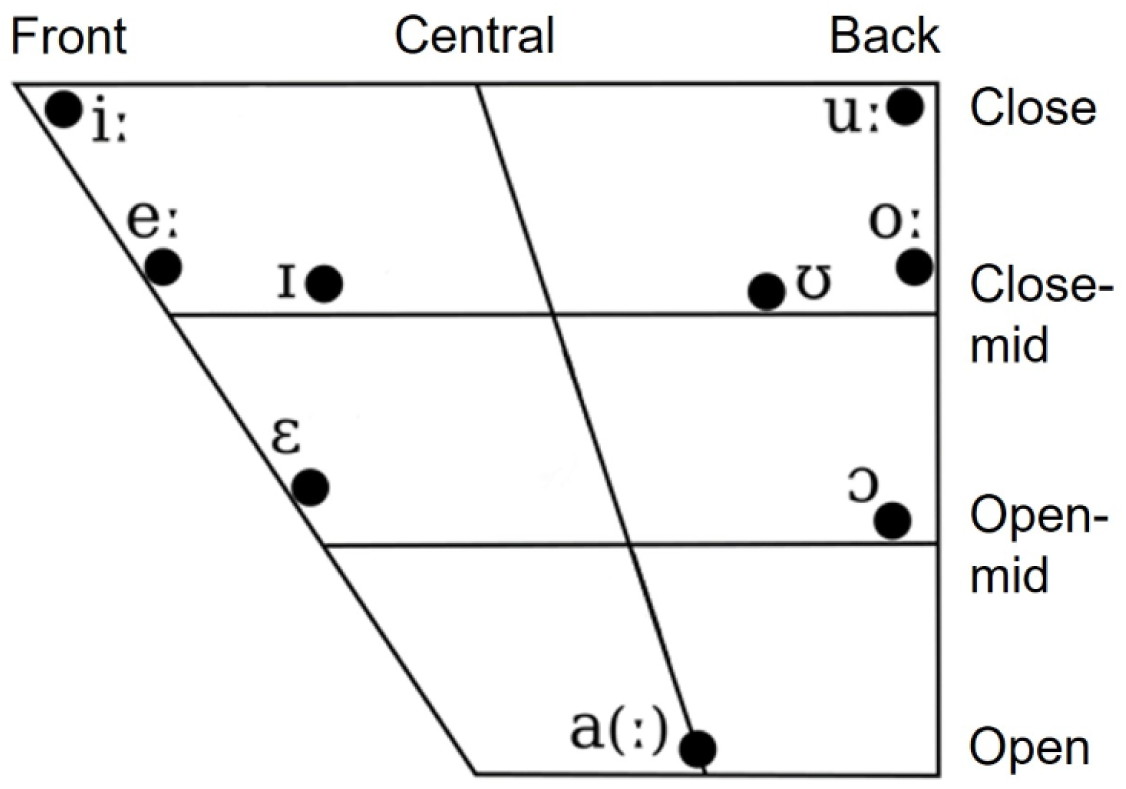
Edited vowel chart for Northern Standard German [62] showing the articulatory configurations of the vowels in the present study.

**S4 Fig.**
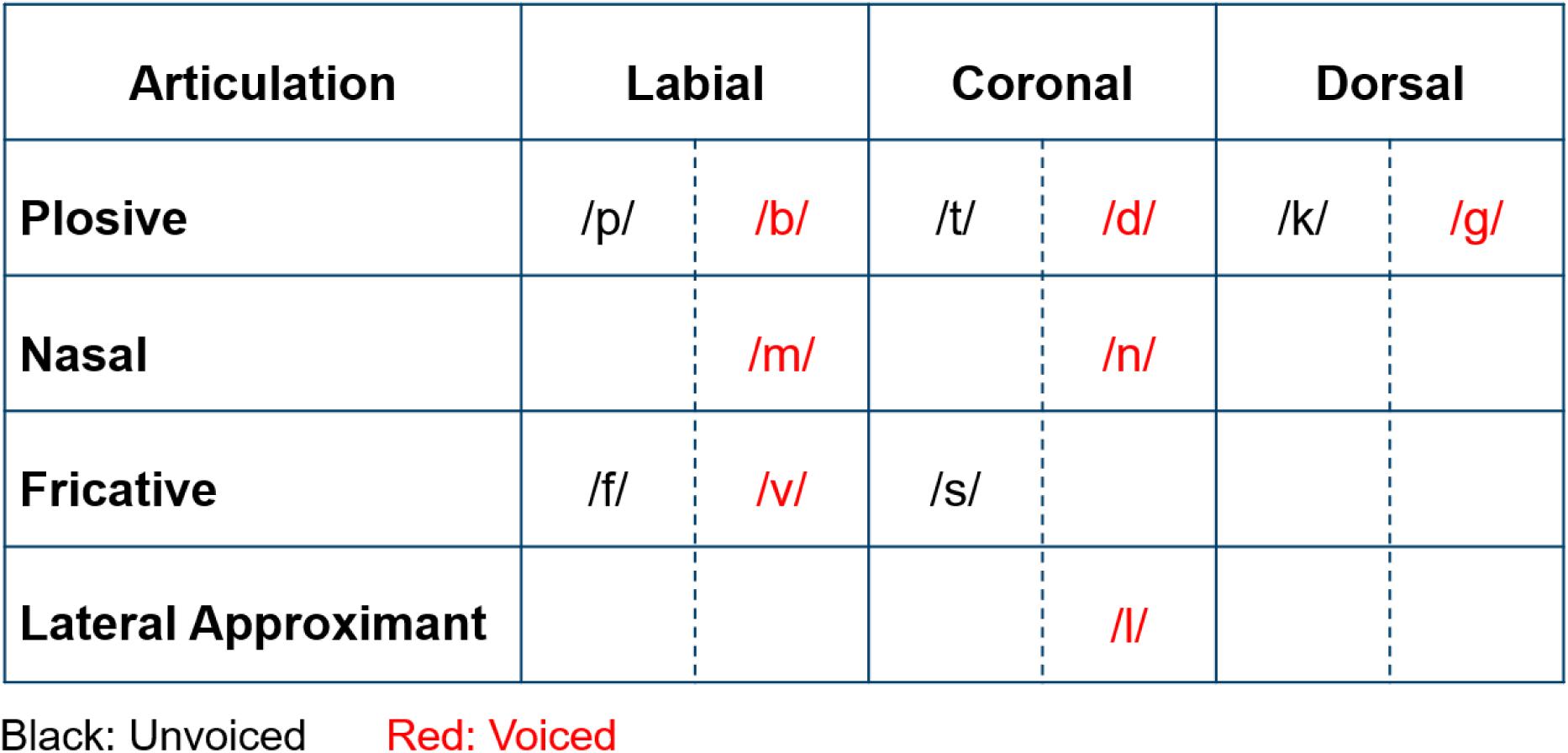
Edited consonant chart [63] showing the articulatory configurations of the consonants in the present study.

**S5 Fig.**
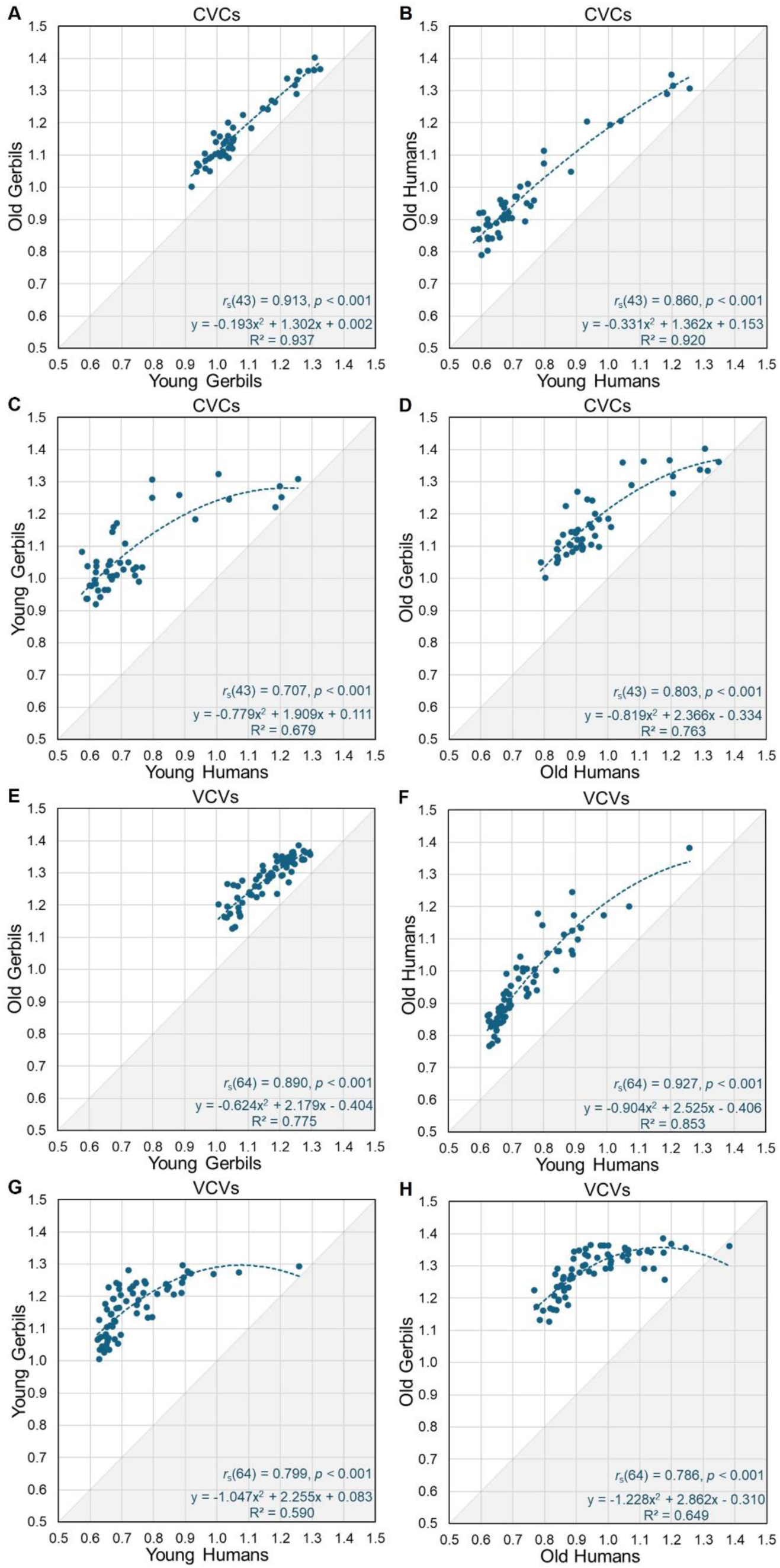
Scatterplots with correlations between mean response latencies for different age groups of gerbils and humans for vowel and consonant discriminations.

## Additional information

### Author contributions

Conceptualization: GMK CJ

Data curation: CJ RB

Formal analysis: CJ

Funding acquisition: GMK

Investigation: CJ CJC

Methodology: GMK CJ RB

Project administration: CJ GMK

Resources: GMK

Software: RB

Supervision: GMK CJ

Validation: CJ RB GMK

Visualization: CJ

Writing – original draft: CJ

Writing – review & editing: CJ GMK RB CJC

### Declaration of competing interest

Declarations of interest: None

### Financial disclosure statement

This work was funded by the Deutsche Forschungsgemeinschaft (DFG, German Research Foundation) under Germany’s Excellence Strategy – EXC 2177/1 - Project ID 390895286).

## Notes

### Competing Interest Statement

The authors have declared no competing interest.

### Summary of Updates

The acknowledgements have been updated to thank the participants of the study.

